# Single cell atlas of trisomy 21 cerebral cortex

**DOI:** 10.1101/2020.01.01.892398

**Authors:** Jingxuan Zhang, Shiyou Wang, Zaoxu Xu, Xiangning Ding, Chen Li, Yonghong Zhang, Yin Chen, Jixing Zhong, Langchao Liang, Chaochao Chai, Xiaoling Wang, Rong Xiang, Jiacheng Zhu, Xiumei Lin, Peiwen Ding, Qiang Zhang, Mingyue Wang, Qikai Feng, Zhijun Zhang, Guangling Guo, Shen Xue, Lin Jin, Zhikai He, Li Yan, Bing Xiao, Changjun Zhang, Yan Xu, Wei Li, Yichi Zhang, Weiying Wu, Sanjie Jiang, Jun Xia, Ya Gao, Lei Wang, Shichen Dong, Si Liu, Shida Zhu, Fang Chen, Dongsheng Chen, Xun Xu

## Abstract

Down syndrome (DS) is one of the most common human birth defects caused by trisomy 21 (T21), leading to a variety of cognitive impairments. The cellular composition of human brain has been explored using single cell RNA sequencing in both physiological and pathological conditions. However, the cellular heterogeneity of human brain with chromosome aneuploidy is largely unknown. Here, we profiled the transcriptome of 36046 cells in cerebral cortex of T21 human fetus, covering frontal lobe, parietal lobe, occipital lobe and temporal lobe. Intriguingly, we detected several genes positively associated with neurons maturation was dysregulated in T21 frontal cortex (*HIC2, POU2F2, ZGLP1* and *FOXK1*). To share, explore and utilized the data resources of T21 cerebral cortex, we developed a comprehensive platform named T21atlas, composing of two functional modules (T21cluster and T21talk). Overall, our study provides, as far as we know, the first single cell atlas for T21 cerebral cortex, which could promote our understanding of the molecular mechanism of DS at an unprecedented resolution and could potentially facilitate the development of novel clinical therapeutics against T21.

## Introduction

Down syndrome (DS), due to the trisomy of human chromosome 21 (Hsa21), was estimated to occur in the frequency of one in every 700 newborns around the world^1^. Due to the presence of an additional copy of Hsa21 compromising ∼250 genes, DS patients demonstrate a wide spectrum of abnormal phenotypes, including early onset Alzheimer disease, early ageing, congenital cardiac defects and childhood leukemia^1^. T21 leads to the degeneration of the cortical pyramidal neurons, deep dendrites, synaptic abnormalities and a reduction of cerebral cortical neurons^2^.

scRNAseq has been widely used to explore the heterogeneity of human cerebral cortex for several years^3-6^. Fan et al. reported the transcriptome of cells from 22 brain regions, resulting in 29 cell sub-clusters with a variety of cells from each region. Zhong et al. obtained the single cell transcriptomes of the developing human pre-frontal cortex ranging from gestation week 8 to 26, from which they identified six cell types composed of 35 sub-types. Through comprehensive analysis, they revealed the intrinsic developmental signals regulating circuit formation and neuron generation^7^. Additionally, brain cellular compositions under pathological conditions have also been studied using scRNA including PD, AD and ASD, revealing cell type and molecular alteration related to pathology^8-11^.

Encouraged by the potent power of scRNA to dissect neural cell diversity as well as cellular micro-environment and its successful applications to understand the development of neural diseases at high resolution and throughput, we performed scRNAseq for four distinct regions of fetal cerebral cortex at mid-gestation period, giving rise to the transcriptome data of 36046 cells. We studied the cellular heterogeneity using unbiased clustering. By comparing our T21 data with previously reported normal brain data^7,12^, we identified dys-regulation of molecular signatures associated with several cell types.

## Results

### Generation of T21 cerebral cortex single cell atlas

To characterize the molecular features of cells in the cerebral cortex in DS brain, we profiled the transcriptome of 36,046 single cells from frontal cortex (FC), temporal cortex (TC), parietal cortex (PC) and occipital cortex (OC) around gestational weeks 23 (Figure 1a). To classify the major cell types, we performed unsupervised cellular clustering using Seurat^13^ to identify transcriptionally similar cells, resulting in seven cell types: excitatory neurons (EX), inhibitory neurons (IN), neural progenitor cells (NPC), oligodendrocyte precursor cells (OPC), astrocytes (AST), endothelia (EN) and microglia (MG) (Figure 1b).

**Figure 1.**
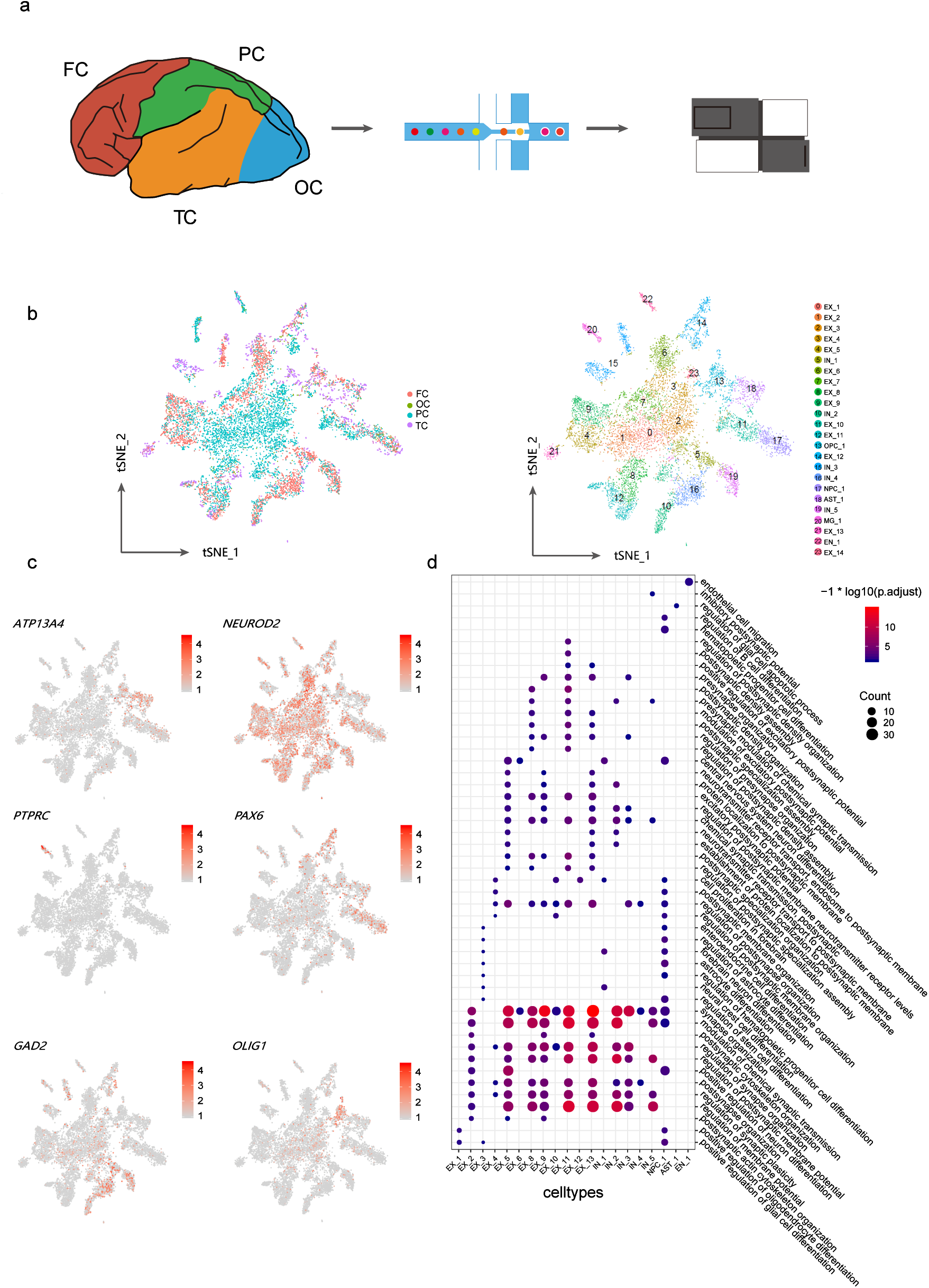

EX highly express *RBFOX1, SATB2* and *NEUROD2* (Figure 1c). *RBFOX1* regulates transcripts TFs alternative splicing, including several critical neural transcripts^14^ and controls neurons excitation. *SATB2* is a TF critical for projection neurons identity and axons function^15^. *NEUROD2* is a transcriptional regulator involved in neuronal determination, highly expressed in mature excitatory neurons. IN specifically expressed canonical markers such as *NXPH1* and *GAD2* (Figure 1c). *NXPH1* contribute to the configuration of *GABA* receptor and synaptic short-term plasticity^16^. *GAD2* is a typical marker of mature inhibitory neurons, catalyzing the production of GABA^17^. NPC specifically expressed *PAX6, TNC, HOPX, SLC1A3* and *GLI3* (Figure 1c). OPC highly expressed *OLIG1*, promoting the formation and maturation of oligodendrocytes^18^ (Figure 1c). AST were indicated by the high expression of *GFAP*^*19*^. MG highly expressed *PTPRC*, encoding CD45 protein associated with immune function (Figure 1c). EN highly expressed typical markers *IGFBP7* and *COL4A2*^*20*^. Besides, we performed Gene Ontology (GO) enrichment analysis based on cell type specific genes. As expected, EX specific genes were enriched in GO terms associated with “synapse organization” and “regulation of synaptic plasticity”. IN specific genes were enriched in GO terms like “regulation of membrane potential”, “chemical synaptic transmission”, “postsynaptic”. NPC specific genes were enriched in GO terms associated with proliferation and differentiation. EN enriched GO terms were related to “regulation of vasculature development” and “endothelial cell migration” (Figure 1d).

### Cellular heterogeneity within T21 cerebral cortex

To further explore the molecular signatures of different cerebral regions, we analyzed the data on the basis of its spatial origin. Briefly, 9326, 4552, 21359 and 899 cells were derived from FC, TC, PC and OC respectively.

Seven major cell types (EX, IN, NPC, OPC, AST, MG and EN), composing of 20 clusters, were profiled from frontal cortex (FC) (Figure 2a). Specifically, excitatory neurons (EX) are composed of 10 clusters which can be grouped into two classes: FC_EX_1, FC_EX_2, FC_EX_3, FC_EX_4, FC_EX_5, FC_EX_6 and FC_EX_7 highly expressing *BFOX1* and *NEUROD2* and FC_EX_8, FC_EX_9, FC_EX_10 specifically expressing *SATB2* and *CUX2* (Figure 2e). Each cluster was discriminated by a subset of unique molecular signatures (Figure 2b). In particular, *DSCAM* was highly expressed in FC_EX_2 (Figure 2d). Previous research showed *DSCAM* was involved in human central and peripheral nervous system development, and this gene was a candidate for Down syndrome^21^.

**Figure 2.**
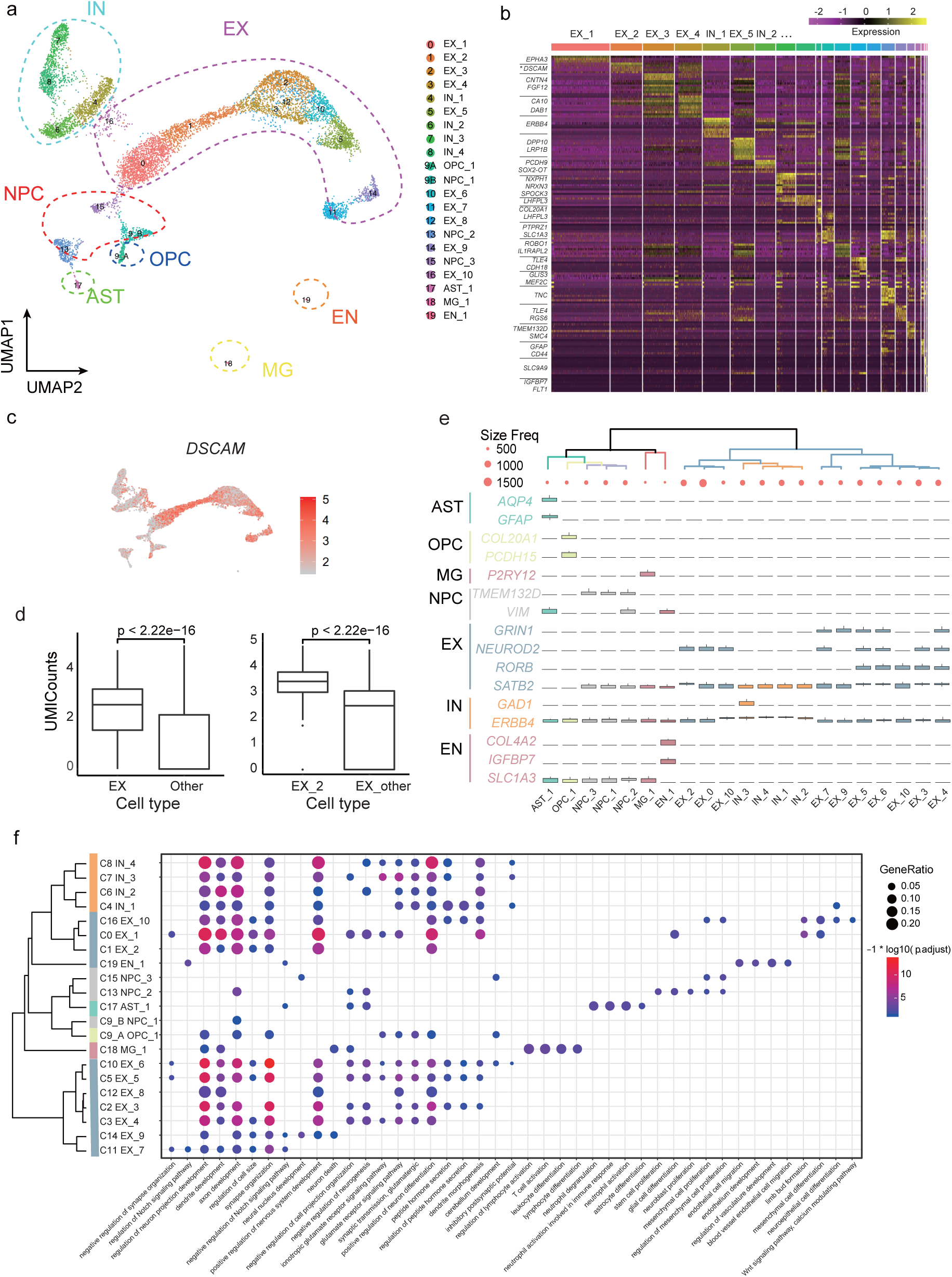

Likewise, FC inhibitory neurons (IN) can be classified into two groups: FC_IN_1, FC_IN_2 expressing *GAD1* and *NXPH1* and FC_IN_3, FC_IN_4 expressing *ERBB4* and *DLX6-AS1*^*4*^. NPC is consisted of three subtypes: FC_NPC_1 highly expressing *PAX6*, FC_NPC_2 and FC_NPC_3 highly expressing *SLC1A3*. FC_AST, FC_MG and FC_EN were composed of one cluster each. Briefly, AST highly expressed *GFAP*. MG highly expressed *PTPRC*, and enriched immune associated GO terms including “regulation of lymphocyte activation” and “T cell activation”. EN highly expressed *IGFBP7* and *COL4A2* and enriched GO terms related to “epithelial cell migration” and “endothelial cell differentiation” associated terms (Figure 2f).

**Temporal cortex (TC)** contribute to 4552 cells (21 clusters) containing six neural cell types (IN, EX, NPC, OPC, AST and MG) and three non-neural cells: endothelial cells (EN), blood cells (BL) and immune cells (IM) (Figure S1). Two clusters (C10, C12), highly expressing *SATB2* and *NEUROD2*, were considered to be mature excitatory neurons. Cluster 2 and 3 (highly expressing *SATB2*) and cluster 13 expressing *SATB2* and *ZEB1* were also annotated as excitatory neurons. Cluster 14 and 18 expressing *NXPH1* and *GAD1*. 21359 cells were retrieved from **parietal cortex (PC)**, from which 18 clusters associated with seven main cell types (IN, EX, NPC, AST, MG and EN) were identified. **Occipital cortex (OC)** data set is consisting of 899 cells which were subsequently classified into nine clusters related to four main cell types (NPC, EX, IN, EN).

### Two group of FC neural progenitor cells with distinct differential potentials and molecular characteristics

To dissect developmental trajectory within FC, we constructed developmental trajectories across the whole frontal cortex. Three trajectories were identified using Monocle3^22-25^ (Figure 3a), with the first trajectory (trajectory 1, hereafter termed as T1) mainly composed of T1_NPC (mainly derived from NPC_1 and NPC_2), OPC and AST while the other trajectory (trajectory 2, hereafter termed as T2) mostly composing of T2_NPC (mostly come from NPC_3) and EX. The third trajectory were mainly composed of IN. This observation is in consistent with previous model that NPC can be differentiated into three lineages: AST, OLG and neurons ^26,27^. In both trajectories, we noticed that NPC were mainly distributed at the root of the developmental path (Figure 3a). We assumed that T1_NPC and T2_NPC may correspond to neural progenitor cells which have the potential to differentiate into OPC/AST lineages and neuron lineage respectively. To reveal the molecular differences between T1_NPC and T2_NPC, we performed differential gene expression analysis between the two NPC groups. We observed that, genes associated with myelination, were enriched in the T1_NPC, such as *QKI* which regulates the differentiation of oligodendrocytes and regulates the myelinating cells of the central nervous system (Figure 3b). We found *HES6* preferentially expressed in T2_NPC (Figure 3b), it has been shown to promote cell differentiation in cortical neurogenesis by inhibiting Hes1 transcription activity in mouse. We found several GO terms specific to each NPC subgroup. Briefly, “myelination”, “ensheathment of neurons” and “axon ensheathment” were specific to T1_NPC while “nuclear division”, “organelle fission” and “chromosome segregation” were unique to T2_NPC (Figure 3c).

**Figure 3.**
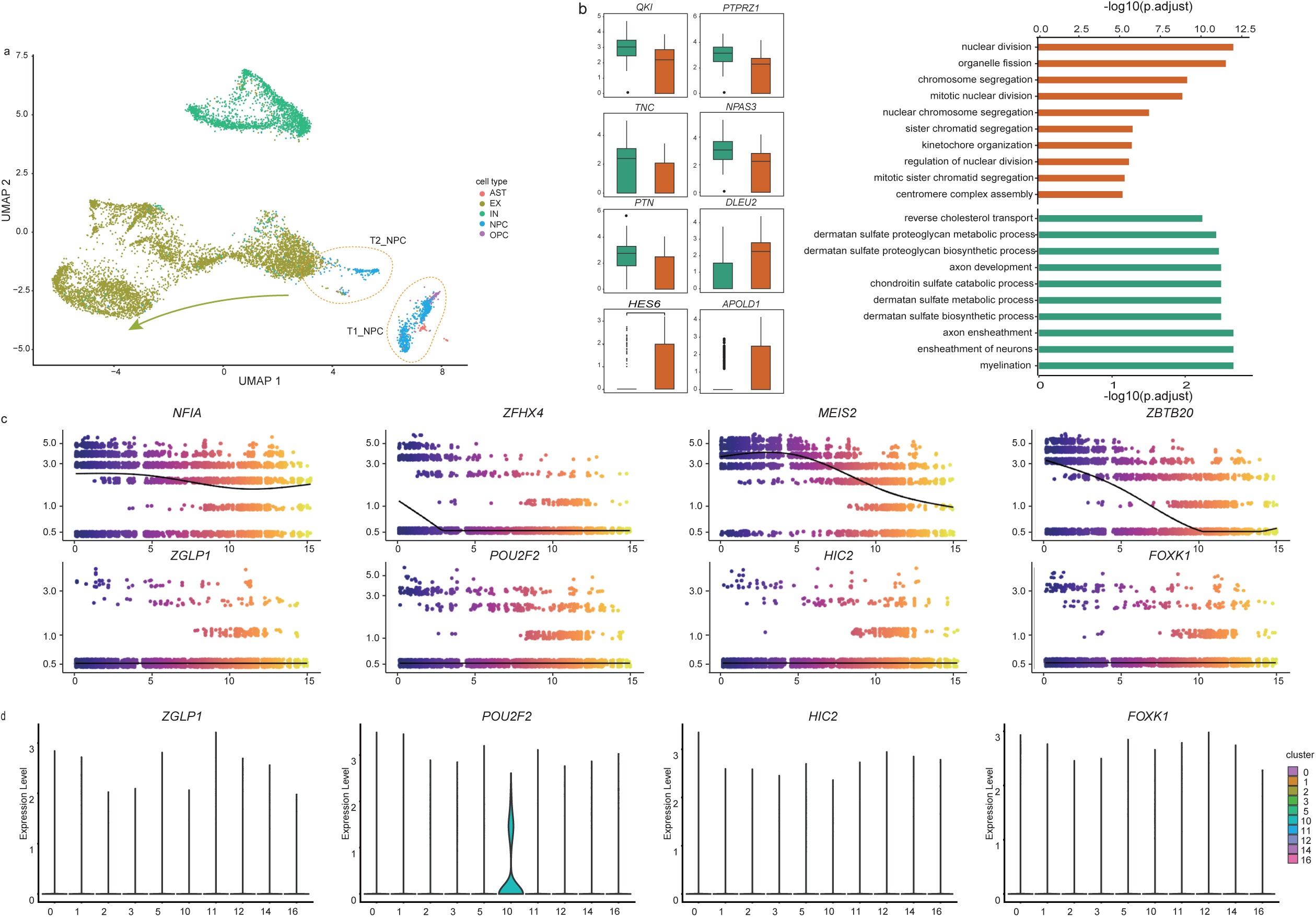

### FC excitatory neuron maturation regulation

Next, we investigated genes associated with FC excitatory neurons maturation. In consistent with previous study, *ZEB1*, the marker of immaturity, declined during the process of EX maturation. Additionally, we found *NFIA, ZFHX4, MEIS2, ZBTB20* were down regulated along excitatory neuron maturation (Figure 3d). This was consistent with the previously reported expression patterns of those genes in normal FC excitatory neurons^7^. According to Fan et al., *ZGLP1, POU2F2, HIC2* and *FOXK1* were up regulated along excitatory neurons maturation process. However, they were expressed in trisomy 21 excitatory neurons at very low levels (Figure 3d), suggesting that the maturation procedure of T21 FC excitatory neurons was distinct from that of EX under physiological condition.

In addition to above mentioned well-characterized neuron maturation regulators, we also identified several putative novel maturation markers (*RORB, TSHZ3* and *SOX5*) in DS excitatory neurons. *RORB* (regulating barrel formation^28^) and *TSHZ3* (playing essential roles in cortical projection neurons development^29^) were up-regulated along FC excitatory neuron maturation trajectory. These up-regulated TFs might be new excitatory neuron maturation markers of DS frontal cortex. In contrast, *SOX5*, was down regulated along excitatory neuron maturation process, implying its role in inhibiting neuron maturation (Figure S3b).

### Cellular communication network among cell subpopulations within cerebral cortex

Cell clusters were reported to express a distinct set of ligands and receptors, which lays the foundation for cell-cell communications. Here, we predicted the putative interactions within FC clusters based on cluster specific ligands/receptors and known ligand-receptor interactions (Figure 4a). Briefly, neural lineage clusters specific ligands range from 10 in cluster EX_1 to 31 in cluster EX_6, and neural lineage clusters specific receptors range from to 17 in cluster EX_1 to 63 in cluster NPC_2. Communications happened most frequently in EX_6, IN_3 and NPC_2, suggesting that several subtypes of neural lineage cells have more frequent interactions than other cell types. In contrast, only few interactions were observed in EX_1, suggesting that cells were rather isolated compared to other neural cell clusters (Figure 4b). Within neural progenitor cells, we further identified 24 interactions were shared by all clusters while cluster NPC_2 was specific to other clusters (Figure 4a). Of particular interest, several interactions have been reported to be important players of neuron function (table s11). For example, *EFNB2* and *EPHA4*, (specifically expressed in EX_2), were proposed that *EPHA4* and its *EFN* binding pairs are expressed by subtype of neuroblasts suggesting the involvement of heterotypic cellular communications and EphA4 signaling functions in NPC niche organization and neuroblast migration in the forebrain in mice ^30^. Besides, our study suggests ligands/receptors were expressed in a cluster specific manner (Figure 4e), *SEMA3A* and its binding partner Plexins such as PLXNA1, PLXNA2 and PLXNA4 exclusively were identified in NPC_2, which could be used to infer the identity of cell clusters.

**Figure 4.**
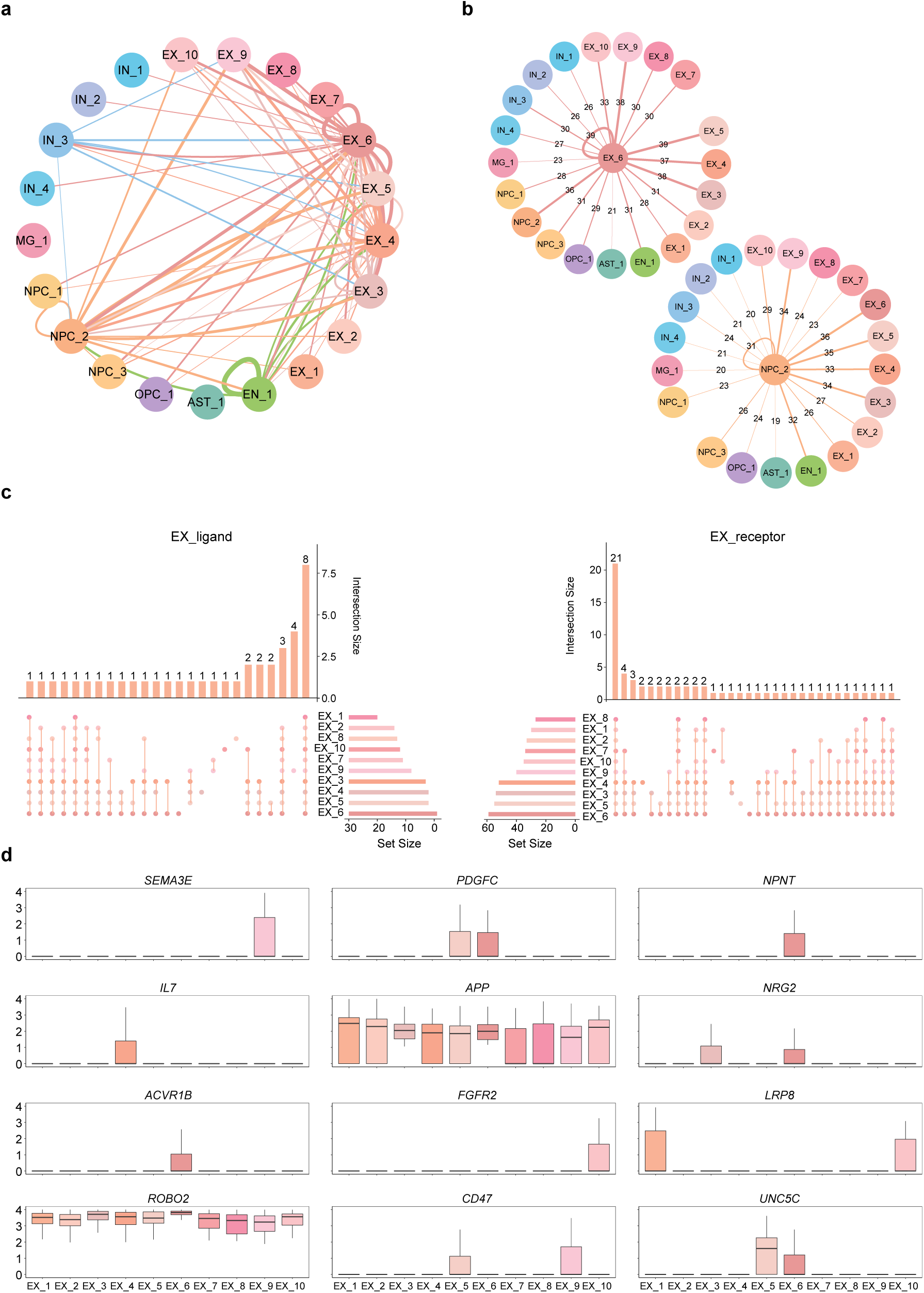

### Genetic regulatory networks of major cell groups in cerebral cortex

Given that the brain is the most complicated important organ composed of diverse cell types governing by sophisticated genetic regulatory modules. The identification of genetic regulatory networks (GRNs) of brain cell populations might provide us some clues about how TFs adapt to specific cellular environment and regulate downstream targets to execute specific biological functions. Here, we constructed the regulatory networks of excitatory neurons, by using cell type-specific TFs and its predicted target genes (Figure 5b). Briefly, we combined cluster specific TFs of each cell type and considered those TFs as cell type specific regulators (Figure 5b).

**Figure 5.**
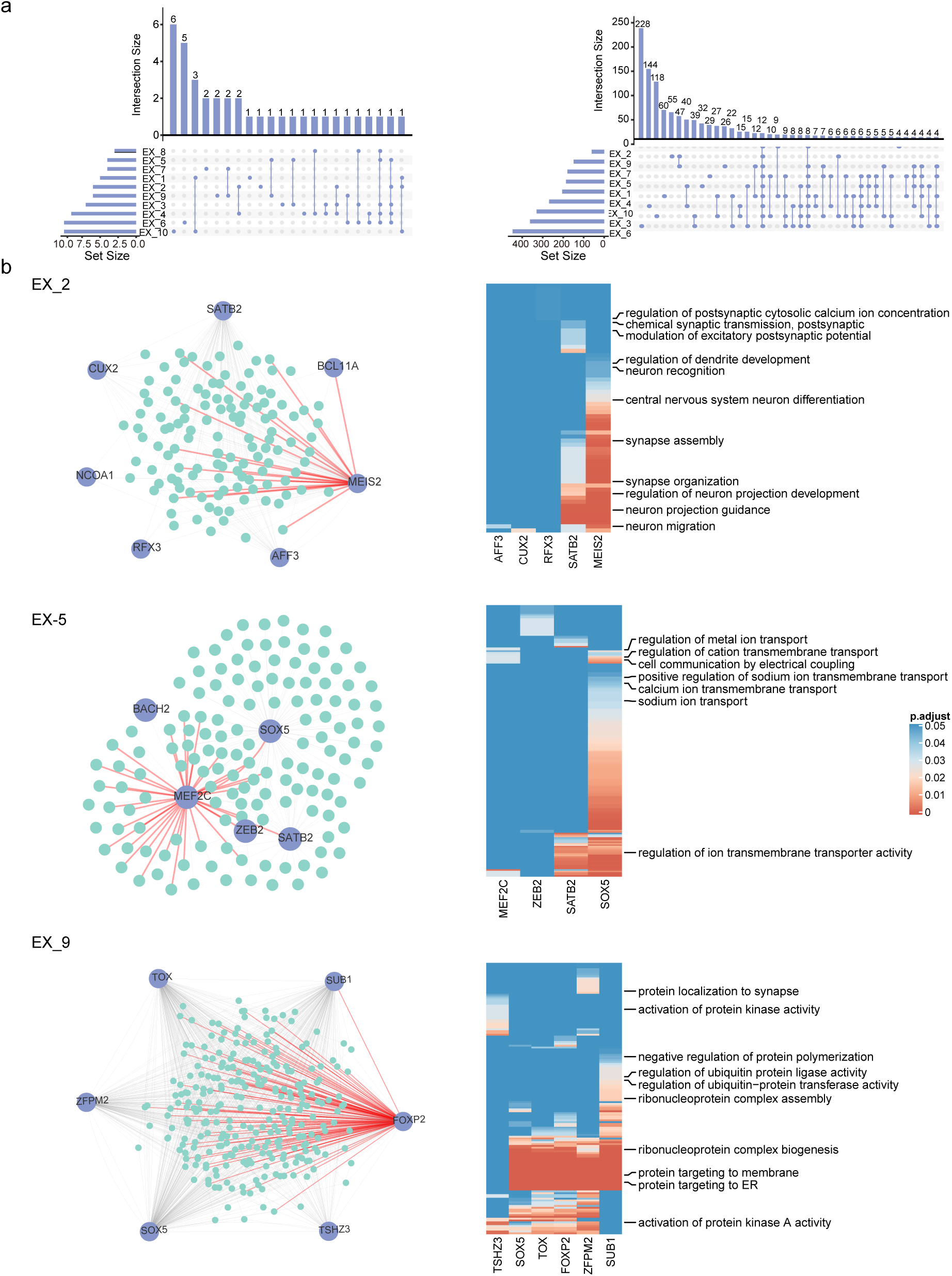

In the EX_2 cell group, six TFs (*MEIS2, BCL11A, SATB2, CUX2, NCOA2* and *RFX3*) were identified as cell type specific TFs. *MEIS2* was regulator of neuron projection guidance and neuron migration ^31^. The GO term of *MEIS2* target genes enriched in axon development, axon guidance and regulation of synapse organization, etc. (Figure 5d). And the TF motif verified targets of *MEIS2* include *DSCAM* which was specifically expressed in EX_2 and was found to be over-expressed in developing CNS ^21^.

In the EX_5 cell group, five TFs (*MEF2C, SOX5, SATB2, XEB2* and *BACH2*) were identified as putative regulators (Figure 5b). Enriched GO terms are mainly related to ion transmembrane transport, cell communication by electrical coupling and modulation of chemical synaptic transmission. Transcriptome analyses revealed MEF2C as putative key regulators in human ASD brains which associated with various genes affected in autism and was a potential therapeutic target for ASD. The MEF2C motif verified target gene (Figure 5e) *ANK3* functions in excessive circuit sensitivity and severe epilepsy by regulating the propagation of action potentials through the assemble of sodium gated ion channels, and found expressed in DS patients ^32^.

In the EX_9 group, six TFs (*ZFPM2, TSHZ3, FOXP2, SUB1, TOX* and *SOX5*) were identified as cell type specific TFs (Figure 5c). The enriched GO terms suggest a strong connection of secretory protein trafficking. For instance, regulome of *FOXP2* converged to specific GO terms including “translational initiation” and “SRP-dependent co-translational protein targeting to membrane”. Additionally, *FOXP2*, by putative interactions with *GRM5, CALM3, NRG3* etc., also seems to modulate retrograde synaptic signaling, an essential process in the generation, maturation and plasticity of synaptic junctions. Taken together, we suspected that EX_9 may be in a state that actively synthesizing and releasing neurotransmitters.

### T21atlas for the sharing and exploring of T21 cerebral cortex single cell atlas

In order to enhance the availability of our data resources, we presented an online interface T21atlas, which is composed of mainly two functional modules: T21cluster module showing the clustering results and corresponding markers, T21talk module covering the putative interaction networks within distinct cell cluster.

## Discussion

Down syndrome has drawn much attentions from researchers in the field of neuroscience. Cognitive impairment is one of the most typical phenotype of DS patient. Although previous studies have investigated DS brain in bulk level, but such data failed to reveal the intrinsic heterogeneity with brain region. Considering the complexity and brain organ, it is necessary to dissect DS in a higher resolution. In this study, we constructed the first single cell atlas of T21 fetal cerebral cortex consisting of 36,046 single cells derived from frontal cortex, parietal cortex, temporal cortex and occipital cortex, revealing the cellular composition and cell population specific molecular signatures.

Neuron maturation is under deliberate regulation by a repertoire of molecules during cortex development. Notably, we found different neural cell distribution in T21 FC and abnormal TFs expression variation in neurons’ maturation. Through trajectory analysis, we identified known neuron maturation markers showing consistent (ZEB1, *NFIA, ZFHX4, MEIS2, ZBTB20*) and altered (*HIC2, POU2F2, ZGLP1* and *FOXK1*) expression patterns in T21 FC. *HIC2* is a key TF encoding gene involved in cytokine-cytokine receptor interaction^38^. *ZGLP1* is a repressor of *GATA* family playing important roles in regulating cell growth^39^. *FOXK1* is reported as the key regulator of neuron maturation^7^. *POU2F2* is a bifunctional regulator repressed or induced neuron differentiation. These dys-regulated TFs might be related to abnormalities of neurons and CNS. Also, we detected several neural markers (*RORB* and *TSHZ3*) which has not been characterized before. We assumed that these up-regulated TFs might represent new excitatory neuron maturation markers controlling FC neuron development.

A distinctive set of TFs were identified from each cell type, representing master regulators coordinating the expression patterns of hundreds downstream targets. To dissect the genetic regulatory network of those cell type TFs, we predicted the targets of each TF based on co-expression patterns. The proliferation, lineage commitment, maturation, reprogramming and apoptosis of a certain cell type is largely determined by several TFs and their genetic regulatory network. Thus, dissecting the regulatory circuits is of crucial importance to understand organism and tissue development. Here, we inferred the targets of TFs based on the Pearson correlation co-efficiency of TF against genes based on the rationale that genes within the same expression module tend to be co-regulated. As each cell types are composed of hundreds to thousands of cells. The genes co-varying at such a sample sale would be very likely to be correlated. And the correlation at single cell level are expecting to reveal TF-target interaction at a sensitive and high-resolution manner. As expected, we predicted a significant proportion of targets for each TF. Among those putative targets, some are well known neural developmental genes.

The single cell resolution defined the characteristics of Down syndrome in a more accurate and sensitive manner compared to traditional methods such as pathology, histology, cell type maker staining and bulk transcriptome, genomic or epigenetic study. Cells are the basic unit of biological functions, exploring the composition of DS brain is of fundamental importance for understanding any degenerative disease. Thus, our data could hopefully serve as the taxonomy of DS cerebral cortex and lays the foundation for future study of DS our findings revealed new insights for investigating the mechanism behind neurologic disease and exploring potential therapies.

## Material and methods

### Ethical statement

Samples were collected with the approval of Medical Ethics Committee of Shiyan Taihe Hospital (201813) following the Code of Ethics of the World Medical Association.

### DS brain collection and dissection

Down syndrome samples collection and dissection was performed the same as previously described^40^. Cerebral cortex dissected from 23 GW fetal brain were separated into small pieces and quick frozen in liquid nitrogen and stored in −80 freezer.

### Single cell RNA sequencing library preparation and sequencing

Nuclei were extracted from frozen brain tissues following previous method. Single nuclei RNA sequencing libraries were constructed using 10X Genomics kit (Chromium™ Single Cell 3’v2) following manufacturer’s instruction. Details of library preparation and sequencing were consistent with methods previously reported^40^.

### Processing of raw single-nucleus RNA-seq data

Raw sequencing data was processed by Cell Ranger 3.0.2 (10X Genomics). Then, Seurat 2.3.4^41^ was applied for downstream analysis. Furthermore, we filtered the data using following criteria: (1) Genes with the UMI under 3 were discarded; (2) Cells with mitochondrial genes percentage greater than 10% were filtered out; (3) Cells with genes number more than 2500 or less than 200 were discarded.

### Cell clustering

Seurat package (v2.3.4) were applied to identify major cell types. Briefly, we used the function FindVariableGenes in Seurat to select the highly variable genes (HVGs) based on the normalized expression matrix (log (UMI/sum (UMI)*10000)). HVGs whose average expression greater than 0.5 and dispersion between 0.0125 and 3 were used as inputs for the downstream analysis. Then we used the function FindClusters in Seurat, to identify cell clusters based on HVGs (for details, see http://satijalab.org/seurat/pbmc-tutorial.html).

### Cluster specific gene identification and GO enrichment analysis

The function FindAllMarkers built-in Seurat were used to identify cluster-specific marker genes (thresh.use = 0.25, min.pct = 0.25, only.pos = TRUE). R package clusterProfiler (10.1089/omi.2011.0118) were employed for the GO term enrichment of cluster specific genes and BH method was used for multiple test correction. GO terms and KEGG terms with P values less than 0.05 were considered as significantly enriched. As for *DSCAM*, Student t test analysis of its expression levels between excitatory neurons and other clusters was performed. In addition, expression levels between cluster 1 (EX) and other EX clusters was examined using t test.

### Developmental trajectory re-construction

Monocle ^42^ was used for developmental trajectory construction based on the set of variable genes, which were selected using the Seurat R package ^41^, with default parameters except we specified reduction_method = “DDRTree” in the reduce dimension function.

### TFs target gene prediction

GENIE3^43^ was used to predict putative target genes of each TF. Then significant target genes for network construction by cytoscape^56^. And targets with transcription factor motifs present in candidate promoter regions were highlighted. Cytoscape 3.6.0^44^ was used to visualize the regulation.

### Cellular communication network

Receptor-ligand relationships were downloaded from IUPHAR/BPS Guide to PHARMACOLOGY ^45^ and Database of Ligand-Receptor Partners (DLRP)^40^. “UMI=0.15” was set as the threshold. Cytoscape was used for the visualization of interactions.

## Data and software availability

This data reported in this study are available in the CNGB Nucleotide Sequence Archive (CNSA: https://db.cngb.org/cnsa; accession number CNP0000816 and CNP0000442)

## Funding

D.C. is supported by China Postdoctoral Science Foundation (grant 2017M622795). This research is funded by the Health Commission of Hubei Province Scientific Research Project (WJ2019M051, ZY2019Q004), the Natural Science Foundation of Hubei Provincial Department of Education (Q20182104),the Free Exploration Project of Hubei University of Medicine (FDFR201609), Cultivating Project for Young Scholar at Hubei University of Medicine (2017QDJZR26, 2016QDJZR11, 2016QDJZR14) and Shenzhen Municipal Government of China JCYJ20180703093402288.

## Acknowledgements

We are thankful to the production team of China National Gene Bank, Shenzhen, China. We appreciate the constructive feedbacks from Professor Guoji Guo.

