## Supplementary material for "Single cell atlas of trisomy 21 cerebral cortex": Figure legend

**Figure 1. Clusters and cell types identified in cerebral cortex of human fetus with T21.**

a. The schematic diagram showing the layout of fetus brain and a brief workflow to generate the single cell transcriptome data in 4 brain regions.
b. t-SNE (left) indicating cells from different regions of brain cortex, namely FC, TC, PC and OC. t-SNE (right) showing clusters identified with all single cells and corresponding cell types they belonged. 7 cell types and 23 clusters were identified in total.
c. Feature plot displaying specific marker genes identified for determination of cell types.

d. Enriched GO terms of differential expressed genes in each cluster.

**Figure 2. Cellular heterogeneity in T21 frontal cortex.**

a. UMAP showing clusters identified in T21 frontal cortex. Dashes of different colors surrounding the clusters indicated corresponding cell types.
b. Heatmap displaying expression of the top10 cluster specific genes. some crucial genes such as *DSCAM* were labeled in the left. Bars above the heatmap indicated different cell clusters.
c. Feature plot showing expression of *DSCAM* gene in each cluster.
d. Boxplot (left) compared *DSCAM* gene expression level of EX with other cell types. Boxplot (right) compared expression of cluster 1 with other clusters within EX cell type. *Student t test* was used with P value indicated.
e. Box plots showing expressions of cell type specific marker genes. Distances between clusters were indicated by the dendrogram and cell frequencies were indicated by circle size.
f. Enriched GO terms of marker genes in each cluster, with cell types and distances indicated in the left.

**Figure 3** **FC NPC groups and FC** **excitatory neuron maturation regulation**

a. Developmental trajectories across the whole frontal cortex without EN and MG.

b. Boxplots showing several differentially expressed markers derived from two different trajectories.

c. Difference of enriched GO term between the NPC subgroups, with the green one representing the first trajectory (mainly derived from NPC_1 and NPC_2) and the orange one representing the other trajectory (mainly derived from NPC_3).

d. Dynamic expression patterns of excitatory neuron along subtrajectory of EX.

e. Violin plot showing the expression levels of *ZGLP1*, *POU2F2*, *HIC2* and *FOXK1* at T21 excitatory neurons.

**Figure 4. Communication network of clusters in frontal cortex**

a. Summary of communication network across and within clusters. Nodes filled with colors indicated clusters of specific cell type. Width of edge correlated positively with the number of ligand-receptor pairs between clusters. Only edges with more than 25 ligand-receptor pairs were made visible.
b. Detailed signaling levels with other clusters of EX_6 and NPC_2, with numbers of ligand-receptor pairs indicated on edges.
c. Cluster specific and shared ligands and receptors numbers of EX.
d. Boxplots showing some of cluster specific and shared ligands and receptors of EX.

**Figure 5. Genetic regulatory networks of major cell groups in cerebral cortex**

a. Venn plot (left) showing cluster specific and shared TFs differentially expressed in EX sub-clusters of FC. Venn plot (right) showing cluster specific and shared GO terms enriched by target genes of TFs differentially expressed in EX sub-clusters in frontal cortex.
b. Interaction network of TFs differentially expressed in EX_2, EX_5 and EX_9 and corresponding target genes. Enriched GO terms by target genes of TFs differentially expressed in EX_2, EX_5 and EX_9.
